# A cost analysis of a cancer genetic service model in the UK

**DOI:** 10.1101/027185

**Authors:** Ingrid Slade, Helen Hanson, Angela George, Kelly Kohut, Ann Strydom, Sarah Wordsworth, Nazneen Rahman, MCG Programme

## Abstract

**Background:** Technological advances in DNA sequencing have made gene testing fast and more affordable. Evidence of cost-effectiveness of genetic service models is essential for successful translation, but remain sparse in the literature. In particular there is a lack of cost data related to genetic services.

**Methods:** A detailed micro-costing of 28 pathways relating to breast and/or ovarian cancer and gene testing for the *BRCA1* and *BRCA2* genes (termed ‘BRCA testing’) was carried out. These data were combined with patient-level data from a Royal Marsden Cancer Genetics Service audit during which BRCA testing was offered to individuals at ≥10% risk of having a mutation.

**Results:** The average cost across all pathways was £2,222.68 (range £376.47-£13,531.24). The average pathway cost for a person with cancer was £1897.71 compared to £2,403.22 for a person without cancer. Of the women seen during audit period, 38% were affected with breast and/or ovarian cancer and 62% were unaffected but concerned about their family history.

**Conclusion:** There is considerable variation in the costs of different gene testing pathways. Improved cost-efficiency could be achieved by increasing the proportion of cancer patients tested, because the pathway cost of an unaffected individual in whom testing has already been performed in a relative with cancer is considerably less.

**Acknowledgements:** We acknowledge NHS funding to the Royal Marsden/ICR NIHR Specialist Biomedical Research Centre for Cancer. SW is supported by funding through the NIHR Oxford Biomedical Research Centre. This work was supported by Wellcome Trust Award 098518/Z/12/Z. For MCG programme see www.mcgprogramme.com.

**Conflict of Interest Statement:** There are no conflicts of interests for any author of this paper

## INTRODUCTION

Health care policy initiatives in recent years in the United Kingdom (UK) and elsewhere have recommended that healthcare services integrate advances in genomic technologies and knowledge into clinical practice for the benefit of patients (1-4). The technological advances are increasingly enabling gene testing to be offered via multigene panel, exome or whole genome testing which potentially allows greater throughput of samples and significant economy of scale (5). However, the decision-making process regarding the affordability of the expansion and integration of genomic technologies into health services is impeded by limited clarity of where costs lie in the standard genetic service model(s) and where genomic testing could best fit into these models.

Clinical cancer genetic units offer services to individuals and families with the goal of assisting treatment decisions in patients with a cancer diagnosis and facilitating early cancer detection and cancer prevention for any future cancers for them and their relatives. Mutations of over 100 genes are known to cause an increased risk of cancer and these underlie approximately 3% of cancer overall (6, 7). There is strong evidence that identification of cancer predisposition gene mutations has an impact on diagnosis and management of cancer patients and their families (6-10).

A large proportion of the work of any cancer genetic service is the management of familial breast and ovarian cancer. Germline mutations in *BRCA1* and *BRCA2* (collectively termed ‘BRCA’) underlie a proportion of both these cancers and the most recent guidelines for familial breast cancer in the UK (2013), recommend testing individuals at ≥10% chance of having a mutation (11).

Typically in the UK, and many other countries, patients requiring assessment for cancer gene testing are referred to a cancer genetic service, where application of risk thresholds, such as the 10% threshold for BRCA testing, recommended the National Institute for Health and Care Excellence (NICE) is used to manage resource allocation (11). However, it has been demonstrated that not all those eligible for testing are being referred to cancer genetic services. For example, ∼15% of high-grade serous ovarian cancer is due to germline BRCA mutations and thus are eligible for testing at a 10% risk threshold (12, 13). However, referral of ovarian cancer patients to genetic services is very low, around 7–20% (9, 12, 13). In part, this is because only about half of mutation-positive ovarian cancer patients report a significant family history of cancer (12, 13). Clearly, important opportunities for improved management of ovarian cancer patients and cancer prevention in their relatives are being missed through the existing processes. Moreover, the cancer genetic service delivery model has limited staff numbers and an infrastructure that is not easily adapted to accommodate unmet need, or to address the increasing demand for cancer gene testing (9).

Evidence of effectiveness and cost-effectiveness of service models, including the use of genomic technologies, is essential for policy-making frameworks but remains sparse in the genetics and genomics literature (14). In particular there is lack of cost data, including readily available published reference costs, for genetic services. Micro-costing, also known as bottom-up costing, is a method requiring identification, measurement and valuation of all underlying activities of a service (15). In this paper, we present a full micro-costing, from the healthcare provider perspective, of a cancer genetic service for breast and ovarian cancer within the UK National Healthcare Service (NHS).

Our study was undertaken as part of the Mainstreaming Cancer Genetics (MCG) programme (www.mcgprogramme.com) a translational initiative that is developing the assays, informatics, clinical infrastructure, education and evaluation to allow implementation of cancer gene testing into routine clinical care of cancer patients and their relatives.

## METHODS

To perform the micro-costing, we first mapped out all the possible pathways relating to breast and/or ovarian cancer and BRCA testing that a patient may follow when referred to the Royal Marsden Cancer Genetics Service prior to the implementation of mainstream testing in June 2013. We believe these to be generally representative of most cancer genetic services in UK in 2014. Once completed, the service activities and resources involved in each step of each pathway were defined and the costs for each activity established so that the overall cost of each patient pathway could be calculated based on 2013 costs.

### Pathway Costings

The patient pathways were defined from referral to surveillance management using service protocols and in discussion with the Cancer Genetics Unit at the Royal Marsden NHS Foundation Trust in London (Figures 1 and 2). The management strategy for each patient was as described in the Royal Marsden Cancer Genetics management protocols in use during the audit period (Appendix 1). These are based on offering testing to those with ≥ 10% risk threshold of having a BRCA mutation, in line with the current NICE guidance (11). If a BRCA mutation is not identified, surveillance recommendations are made according to the individual’s family history. Population surveillance recommendation was costed for mammography 3 yearly from 50 to 70 years of age; moderate risk surveillance is annual mammography from 40 to 50 years of age and then entering the population surveillance; higher risk surveillance is annual mammography 40 to 50 years eighteen monthly mammograms from 50 to 60 years and then entering the population surveillance programme as per RMH protocols (Appendix 2) (15).

**Figure 1:**
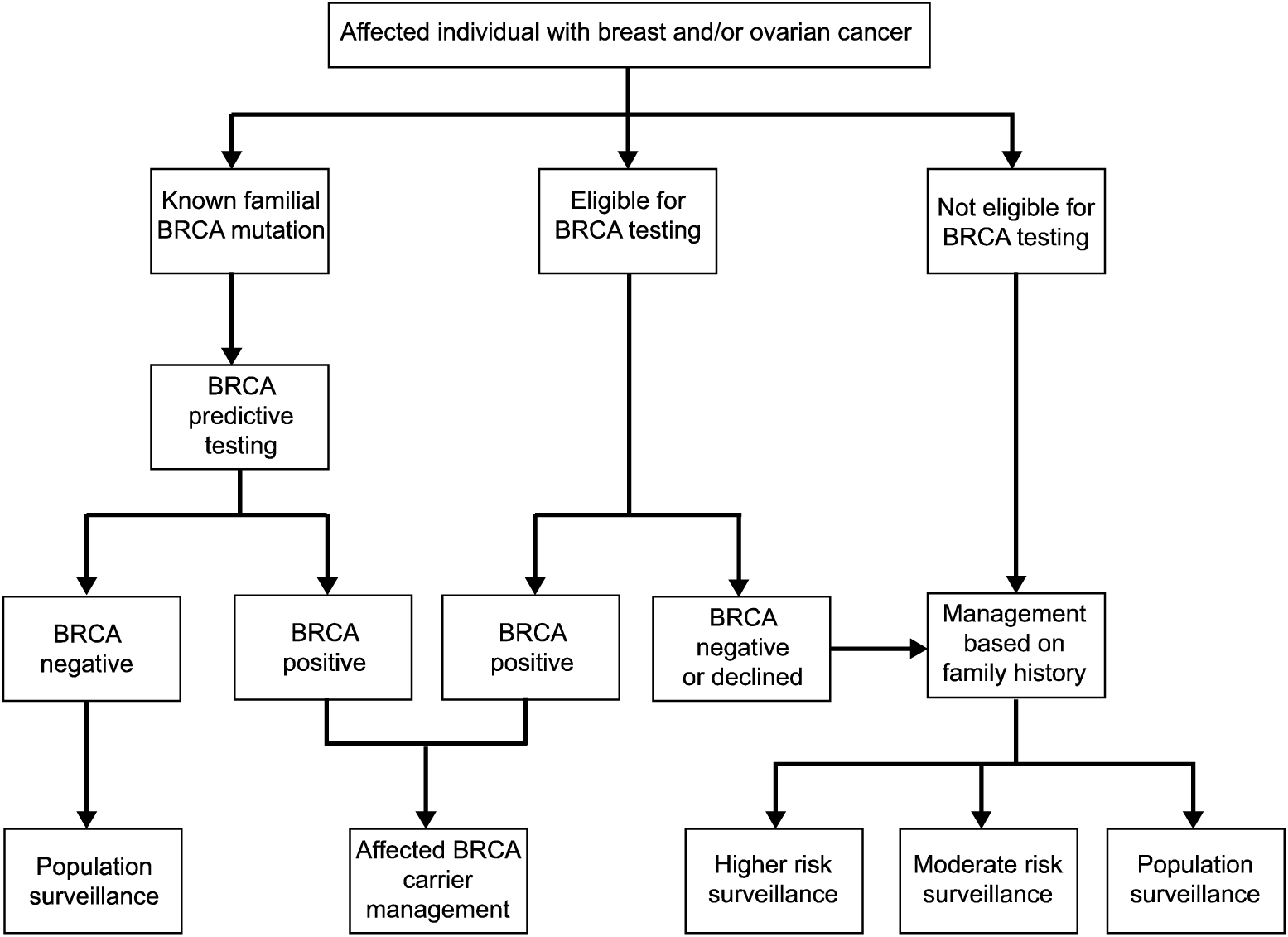
**Individual affected with breast or ovarian cancer – patient pathways**

**Figure 2:**
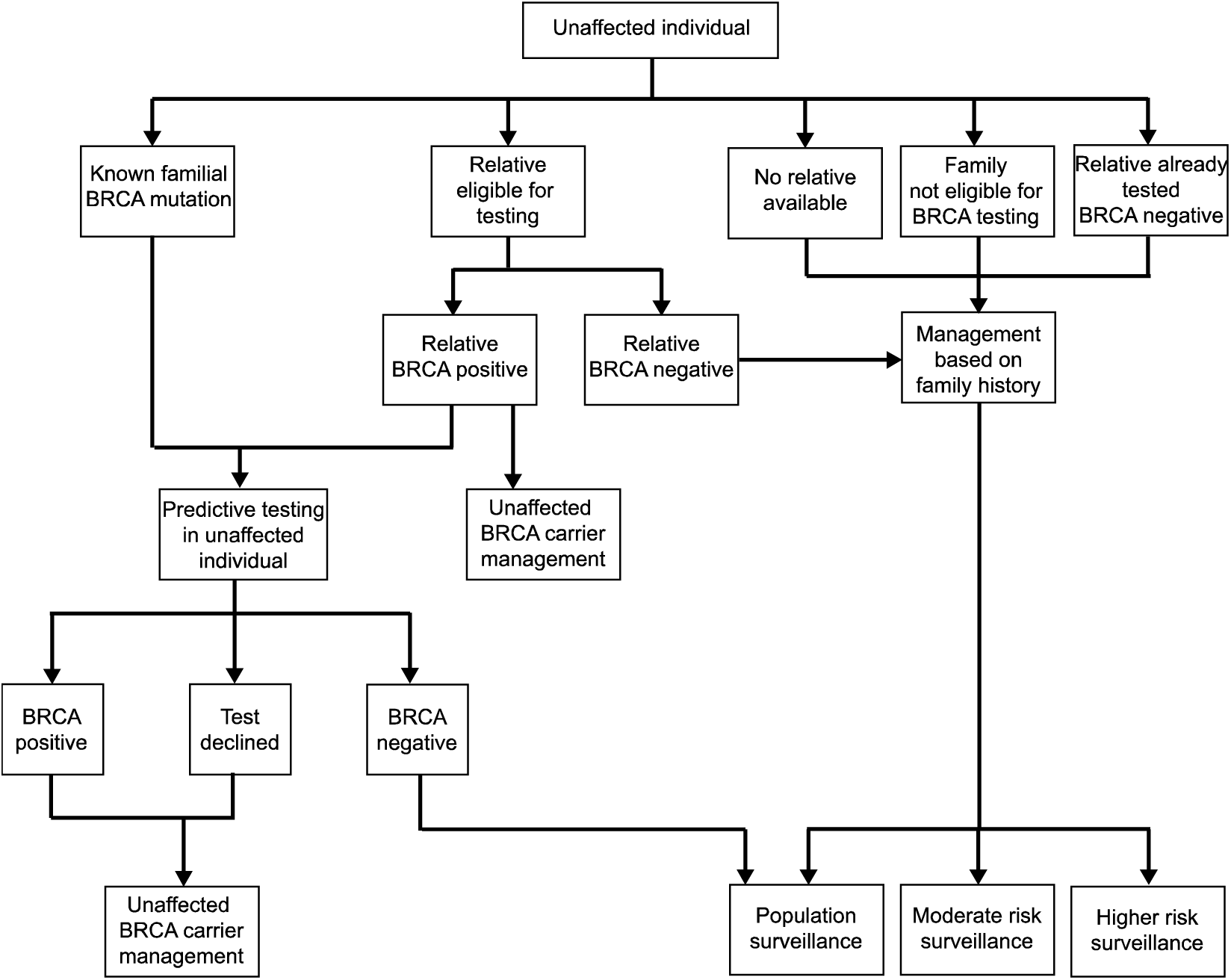
**Individual unaffected - patient pathways**

In order to maintain a manageable number of patient pathways, in families with no BRCA mutation it was assumed that relatives were in the same risk group as the proband. In families where an individual with a BRCA mutation is identified, relatives either carry the mutation (or chose not to be tested) with subsequent surveillance management as mutation carriers, or they do not carry the familial mutation and are managed with population-level surveillance through the NHS. For individuals where a BRCA mutation is identified, carrier surveillance comprises annual mammography 40 to 70 years and annual MRI 30 to 50 years. In addition, mutation carriers are eligible for risk reducing surgery; bilateral mastectomy and/or bilateral salpingo-oophorectomy. The uptake of these interventions in unaffected carriers was determined using expert opinion from the clinical unit alongside literature estimates; 30% for bilateral mastectomy and 60% for bilateral salpingo-oophorectomy (16-18). For affected carriers, the 5 year survival rate at age 40 years (0.7 for breast cancer and 0.69 for ovarian cancer www.ons.gov.uk) was incorporated. It was assumed that 5% of ovarian cancer patients with a BRCA mutation that survive to 5 years would undergo bilateral mastectomy. These rates were based on discussions with the Royal Marsden Cancer Genetics Unit, as no published data were available.

In order to identify resource use in each pathway clinical, administrative and laboratory staff were asked to estimate the length of time each defined activity took them in minutes, the general quantity of consumables, and where each activity took place in the pathway. The full BRCA test includes comprehensive analysis of the full coding sequence and intron-exon boundaries for small intragenic mutations and larger exonic deletion/duplications. A predictive test describes a targeted analysis for a specific mutation already known to predispose to cancer in the family. For this paper we used the TGLclinical Sanger sequencing + MLPA sequencing cost for full gene testing, which was charged to the Royal Marsden NHS Foundation Trust in 2013 exclusive of VAT costs as per NICE guidance. It should be noted that there is variability in BRCA gene test costs across the NHS and the TGLclinical test cost was at the lower end of the range. The NICE guidance used a comparable test cost of £700 which is reflected in the sensitivity analysis (11). Additionally the testing is now performed with NGS technology using the TruSight Cancer Panel (see www.TGLclinical.com for further details). The cost of post genetic testing management, which includes mammography, MRI, mastectomy and salpingo-oopherectomy, was taken from 2013 NHS reference costs which are published average costs derived from hospital trust submissions and the literature (11, 19, 20).

Staff salary unit costs for administrators, laboratory and clinical staff were obtained from either the NHS agenda for change or Personal Social Services Research Unit (PSSRU) reference costs for 2013 (8, 19). The mid-point of each grade was used and National Insurance, superannuation and overhead costs added if not already included. In addition, the cost of genetic counsellor time accounts for supervision at a ratio of one hour consultant supervision to 12.5 hours counsellor time (21). All staff time was calculated with a London weighting of 1.19 as outlined in the PSSRU (reference costs) (Table 1) (22). The costs of clinical appointments were taken from the 2011-2012 NHS reference costs (oncology) and PSSRU costs (general practice) (19, 22).

**Table 1.** **Unit costs for clinical activity and patient management costs (2013)**

|  |  |  |
| --- | --- | --- |
| **Cost Items** | **Cost (£)** | **Source of Data** |
| **Genetic Service Activity** |  |  |
| Referral received and processed | 5.07 | Primary data collection |
| Referral triaged | 1.17 | Primary data collection |
| Request documents | 8.82 | Primary data collection |
| Clinical review of case | 21.85 | Primary data collection |
| Appointment arranged | 3.49 | Primary data collection |
| Clinic preparation | 2.74 | Primary data collection |
| Clinic appointment | 104.80 | Primary data collection |
| Post appointment letter | 11.18 | Primary data collection |
| Post appointment administration | 5.17 | Primary data collection |
| **Staff Salaries (London)** |  |  |
| Band 3, administrative assistant | 17.19 per hour | NHS Agenda for change |
| Band 5, medical secretary | 23.48 per hour | NHS Agenda for change |
| Band 6, administrative lead | 28.74 per hour | NHS Agenda for change |
| Band 8, genetic counsellor | 55.54 per hour | NHS Agenda for change |
| Registrar | 61.46 per hour | PSSRU |
| Consultant | 139.73 per hour | PSSRU |
| **Patient Management** |  |  |
| Mammography | 45.50 | Reference (20) |
| MRI | 145.88 | Reference (19) |
| Mastectomy | 6,784.00 | Reference (11, 19) |
| Salpingo-oophorectomy | 3,355.43 | Reference (11, 19) |

The main (base case) analysis assumes that those affected by cancer were referred through their oncologist, and those unaffected by cancer were referred by their general practitioner. This analysis is restricted to women not within populations in whom BRCA founder mutation testing is available, such as the Ashkenazi Jewish population. Furthermore, the base case analysis assumes that all women are seen in the cancer genetic service at the age of 30 years. This age was chosen as it accommodates the years of highest relative risk of both breast and ovarian cancer in BRCA mutation carriers (23). All costs are presented in 2013 pounds Sterling and assumed to be incurred at the point of service delivery with the exception of surveillance which continues until the age of 70 years and hormone replacement therapy which continues until the age of 50 years. A discount rate of 3.5% has been applied to the costs associated with surveillance and hormone replacement therapy (15). This discounting rate adjusts the costs to reflect both time preference and the fact that items depreciate over time. Risk reducing surgery is assumed to take place in the year of diagnosis and therefore not subject to discounting. (Table 1)

#### Audit Data

An audit of all clinical activity relating to breast and/or ovarian cancer patients or unaffected patients with a family history of breast and/or ovarian cancer was undertaken at the Royal Marsden Cancer Genetics Service between September 2009 and February 2010. These data were used to determine the number of patients, at first appointment, that followed each patient pathway within the audit period. Women, with breast and/or ovarian cancer, whose consultation was in relation to BRCA testing, were included; excluding those tested for founder mutations and other cancer predisposition genes and male patients. This audit excludes those that were inappropriately referred or failed to attend appointments, these additional service considerations were included in the sensitivity analysis.

#### Sensitivity Analysis

In order to examine how sensitive the costing results were likely to be to any assumptions that we made, sensitivity analysis was performed where some elements of resource use and costs were varied. The costs varied in the sensitivity analysis included member of clinical team involved in the patient pathway, method of consultation, proportion of patients who did not attend a first or second appointment, cost of the test and removal of London weighting.

### RESULTS

#### Patient Pathways

A total of 28 individual patient pathways were identified for the delivery of the breast and ovarian cancer genetic services, which are split into individuals affected and unaffected by cancer. Full description of all 28 pathways and their associated costs are included in Supplementary Tables 1 and 2. Pathway 6 is shown in Table 2 as an exemplar, chosen as it is the most frequent pathway for cancer patients. This pathway is for an individual with cancer that is eligible for BRCA testing, who has a negative BRCA test.

**Table 2.** **Example of patient pathway units of activity, and associated costs**

|  |  |  |
| --- | --- | --- |
| Pathway 6 Activity | Sub Activity | Cost (£) |
| Oncology referral | Oncology clinic appointment | 168.00 |
| Appointment administration | Referral received and processed | 5.07 |
|  | Referral triaged | 1.17 |
|  | Request documents | 8.82 |
|  | Clinical review | 21.85 |
|  | Appointment arranged | 3.49 |
|  | Clinic preparation | 2.74 |
| Clinic related activity | Clinic appointment | 104.80 |
|  | Post appointment letter | 11.18 |
|  | Post appointment administration | 5.17 |
| Blood sample | Phlebotomy | 3.00 |
| BRCA full gene test |  | 540.00 |
| Follow up appointment administration | Appointment arranged | 3.49 |
|  | Clinic preparation | 2.74 |
| Follow up clinic related activity | Clinic appointment | 104.80 |
|  | Post appointment letter | 11.18 |
|  | Post appointment administration | 5.17 |
| Higher risk surveillance | Mammography | 628.30 |

The supplementary tables and Table 3 gives a detailed breakdown of these pathways, but in summary, pathways 1 to 10 represent individuals referred to the cancer genetic service, who were affected with breast and/or ovarian cancer (affected patient pathways). The main differences between these 10 pathways are related to whether individuals are considered to be eligible for BRCA testing, whether they decide to undergo testing, and their subsequent management based on the test result and their family history.

**Table 3.**
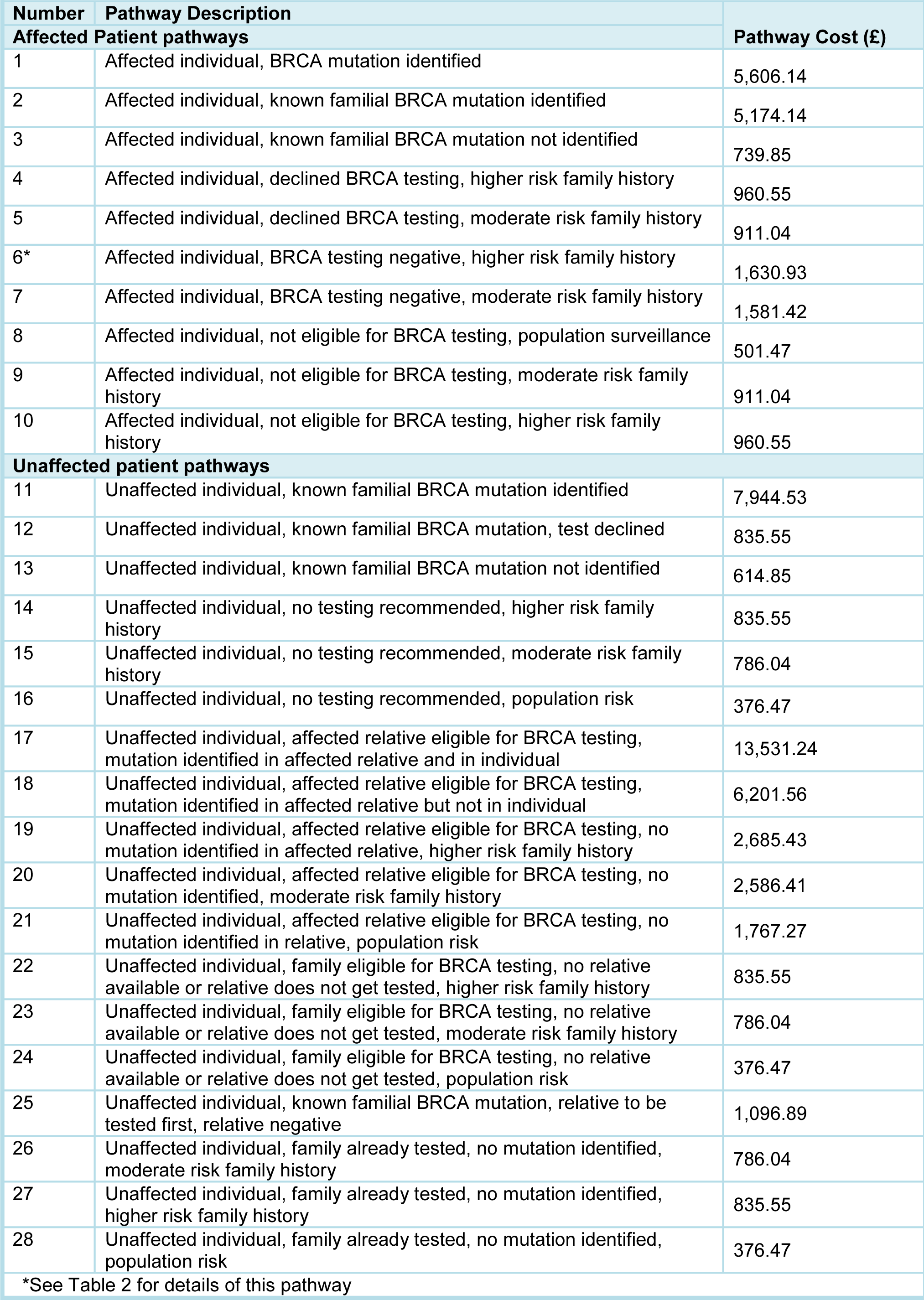
**Pathway costs**

Pathways 11 to 28 represent individuals referred to cancer genetic services, who are not affected with cancer (unaffected patient pathways). The main differences between these pathways are related to whether a relative of the individual has previously undergone BRCA mutation testing, and if not, whether the individual or a relative is eligible for BRCA testing, and their subsequent management based on the test result and their family history.

##### Pathway Costs

Table 3 presents the total cost of each individual pathway. The most expensive pathway (£13,531.24) is that in which an unaffected individual presents, a relative of that individual has a BRCA test which is positive and the unaffected individual is also mutation positive. This reflects the management costs incurred by both individuals in the family. There are three pathways with the lowest cost (£376.47), each of which are pathways for unaffected individuals either from a family where previous testing has not identified a BRCA mutation, from a family where no relative is available for testing or from a family where no BRCA testing is recommended. In each pathway the unaffected individual is subsequently managed at population risk. This lower cost represents the absence of genetic testing and the lower management costs.

The average cost across all 28 patient pathways in the cancer genetic services was £2,222.68 per pathway, the average cost per pathway for a person affected with breast or ovarian cancer was £1897.71 compared to £2403.22 for an unaffected person. The average cost of a pathway where the presenting patient is found to carry a BRCA mutation was £8064.01 representing higher management costs in these patients. The average cost of a pathway where a full BRCA test was carried out in a presenting patient or relative was £4,448.80 compared to £3,114.05 when a predictive BRCA test for a known mutation is performed. In the situation of BRCA testing being recommended in a relative of the presenting unaffected patient, the average cost of the pathway rose to £5,354.38.

##### Audit Data

A total of 220 women had first appointments, regarding breast and/or ovarian and BRCA testing, within the Cancer Genetics Service at the Royal Marsden NHS Foundation Trust, between September 2009 and February 2010. Of these 84 (38%) women were affected with breast and/or ovarian cancer and 136 (62%) were unaffected but concerned about their family history. A total of 72 (33%) women were eligible for BRCA testing at the first appointment either as a full mutation screen (n=42) or as a predictive test (n=30).

Of the 84 women with breast and/or ovarian cancer, seen in first appointments between September 2009 and February 2010, 42% (35/84) were eligible for, and underwent, BRCA testing. In nine patients a mutation was identified. In the 26 where no mutation was identified 23 were eligible for higher risk surveillance and three for moderate risk surveillance. Seven women declined genetic testing, all of whom were eligible for higher risk surveillance. In 32 women no testing was recommended; four were at population risk, 12 were eligible for moderate risk surveillance and 16 for higher risk surveillance. There were a further 10 women affected with breast and/or ovarian cancer, who had not themselves been tested, but where a familial BRCA mutation was known.

A total of 136 unaffected individuals were seen in first appointments during the audit period. In 30/136 cases there was a known familial BRCA mutation, 22 of these underwent predictive testing; seven were found to carry a mutation. There were 35 of these 136 unaffected women in whom BRCA testing in a family member was not recommended; eight were population risk, 18 eligible for moderate risk surveillance and nine higher risk surveillance.

BRCA testing in a relative with cancer was recommended for 19 unaffected individuals. In three of these 19 cases there was a known familial mutation in a distant relative and an intervening relative required testing prior to further management of the patient. In each of these families the intervening relative was found not to carry a BRCA mutation and therefore testing was not carried out in the unaffected patient. In 16 of these 19 cases it was recommended that an affected relative had a full BRCA test; four of these relatives were mutation positive, allowing the unaffected individual to undergo predictive testing. The period of the audit precludes inclusion of results of these predictive tests, a probability of 50% was used to predict that two of these four unaffected individuals would be BRCA mutation carriers and two would not. There were 12 relatives found not to carry a mutation; 11 of these were families eligible for higher risk surveillance and one was at population risk.

For 12 unaffected individuals an affected relative had already been tested and found to be negative for BRCA mutations. Three were eligible for moderate risk surveillance and nine for higher risk surveillance. For 40 unaffected individuals their families were potentially eligible for testing but either no relative with cancer was available for testing, a relative was informed about eligibility but did not have testing or the unaffected individual decided not to proceed with testing. Thirty-two of these unaffected individuals were eligible for higher risk surveillance and five for moderate risk surveillance. Three cases were at population risk.

##### Sensitivity

The results of the sensitivity analysis show that varying the test cost had the greatest impact on total service costs whilst varying the proportion of appointments undertaken by genetic counsellors had the least impact on overall cost of service (Table 4).

**Table 4.** **Sensitivity analysis of total service cost in six month period**

| Table 4. Sensitivity analysis of total service cost in six month period |  |  |
| --- | --- | --- |
| Sensitivity Analysis | Cost Varied | Cost of Service During Audit Period (£) |
| Base Case Cost (100% consultant appointments, 100% face to face appointments, cost of test £540, London weighting) |  | 386,993 |
| Removal of London Weighting |  | 378,352 |
| Varying Test Cost | test at £300 | 374,753 |
|  | test at £400 | 379,853 |
|  | test at £600 | 390,053 |
|  | test at £700 | 395,153 |
|  | test at £1000 | 410,453 |
| Varying Proportion of Consultant Appointments | 25% consultant, 75% genetic counsellor appointments | 377,178 |
|  | 50% consultant, 50% genetic counsellor appointments | 380,450 |
|  | 75% consultant, 25% genetic counsellor appointments | 383,721 |
| Varying Proportion of Clinic Appointments | 25% telephone, 75% clinic appointments | 383,487 |
|  | 50% telephone, 50% clinic appointments | 379,980 |
|  | 75% telephone, 25% clinic appointments | 376,474 |

#### DISCUSSION

The detailed costing analysis presented here provides an insight into the resources required in the delivery of the current cancer genetic service for breast and ovarian cancer. Twenty-eight patient pathways were identified with associated costs ranging from £376.47 to £13,531.24 (difference of £13,154.77) depending on the testing strategy and management plan for the patient.

The burden of cost in the patient pathways presented here lies in the management of patients, in particular those identified as carrying a BRCA mutation. To fully evaluate cost-effectiveness these data would need to be combined with outcome data, for example to include the costs saved from cancers prevented through risk-reducing surgery. The available data suggest that identifying BRCA mutations is cost-effective (11, 24-26). Of particular relevance in the UK, the NICE guidance for familial breast cancer, using economic modelling, determined that testing affected or unaffected individuals at a risk threshold of ≥5% was cost-effective in women under 59 years (at a CE threshold of £30,000), although the clinical guidance recommends testing for women of any age, at a risk threshold of ≥10% (11).

Intuitively, an unaffected individual would be expected to receive the maximum benefits of genetic testing such as a reduced incidence of primary cancers. Furthermore, in individuals found to be mutation negative cost savings are generated from reduced surveillance (11). From the service perspective the most cost-efficient strategy would be to identify unaffected relatives from an affected individual in whom a BRCA mutation has been identified. These pathways are demonstrated to have an average cost of £3,114.05. Moreover, these pathways reduce the time and expense of the ‘loops’ seen when an unaffected individual is referred to the cancer genetic service but, though eligible, their relative with cancer has not been offered BRCA testing. In this scenario, i.e. where BRCA testing is recommended in an affected relative of an unaffected patient seen in genetics, the average pathway cost rises to £5,354.38. Two-thirds of patients referred to the cancer genetic service were unaffected and only one third of these were from a family where BRCA mutation testing had been performed in an affected relative.

One mechanism for reducing the time and expense of these ‘loops’ would be if individuals with breast and/or ovarian cancer eligible for BRCA testing were more routinely getting access to genetic testing. Cancer genetic services have restricted capacity, and would be unlikely to be able to deliver testing comprehensively to cancer patients (9). However, more access to genetic testing could be possible if testing in cancer patients became integrated into oncology services. This service model has been termed the ‘mainstreaming’ of genetic testing (7, 27, 28). The MCG programme, and other initiatives, are developing and currently implementing mainstreaming for BRCA testing through oncology (www.mcgprogramme.com), in close communication with genetics. Even if it is assumed that initial expenditure may increase due to higher volume of tests being undertaken, although new technologies may mitigate against this increase, the identification of more affected individuals would facilitate efficient cascade testing and preventative measures saving overall expenditure for the healthcare provider (11).

Additionally there is potential for this approach to offer two substantial advantages. Firstly, it would deliver greater equity of access to BRCA testing at guideline thresholds, and secondly it would streamline the clinical pathways making them more time-efficient for the patient and the clinical teams, with the capacity to accommodate growing demand. It is likely that the integration of cancer genetic testing into routine patient pathways in oncology, in close liaison with genetic services, will prove to be the optimal pathway for most cancer patients. However, cost-effectiveness analysis is required to better understand the benefits that would be gained from this broadening of testing. Furthermore, the evaluation of new sequencing technologies needs to be built into this analysis to explore potential additional efficiency savings provided by the increased throughput and reduced costs.

Our paper has provided a basis for understanding what resources are currently being used in cancer genetic services, so that policy makers can better understand the starting point for integrating cancer gene testing, and in the future new sequencing technologies, into cancer care. Along with the development of mainstream testing pathways, and in future the harnessing of new sequencing technologies for clinical diagnostics, a comprehensive translational evidence base including service evaluation, economic evidence and careful consideration of the resource allocation challenges are essential to making genomic medicine a reality (14, 29).

---

**Supplementary Table 1:**
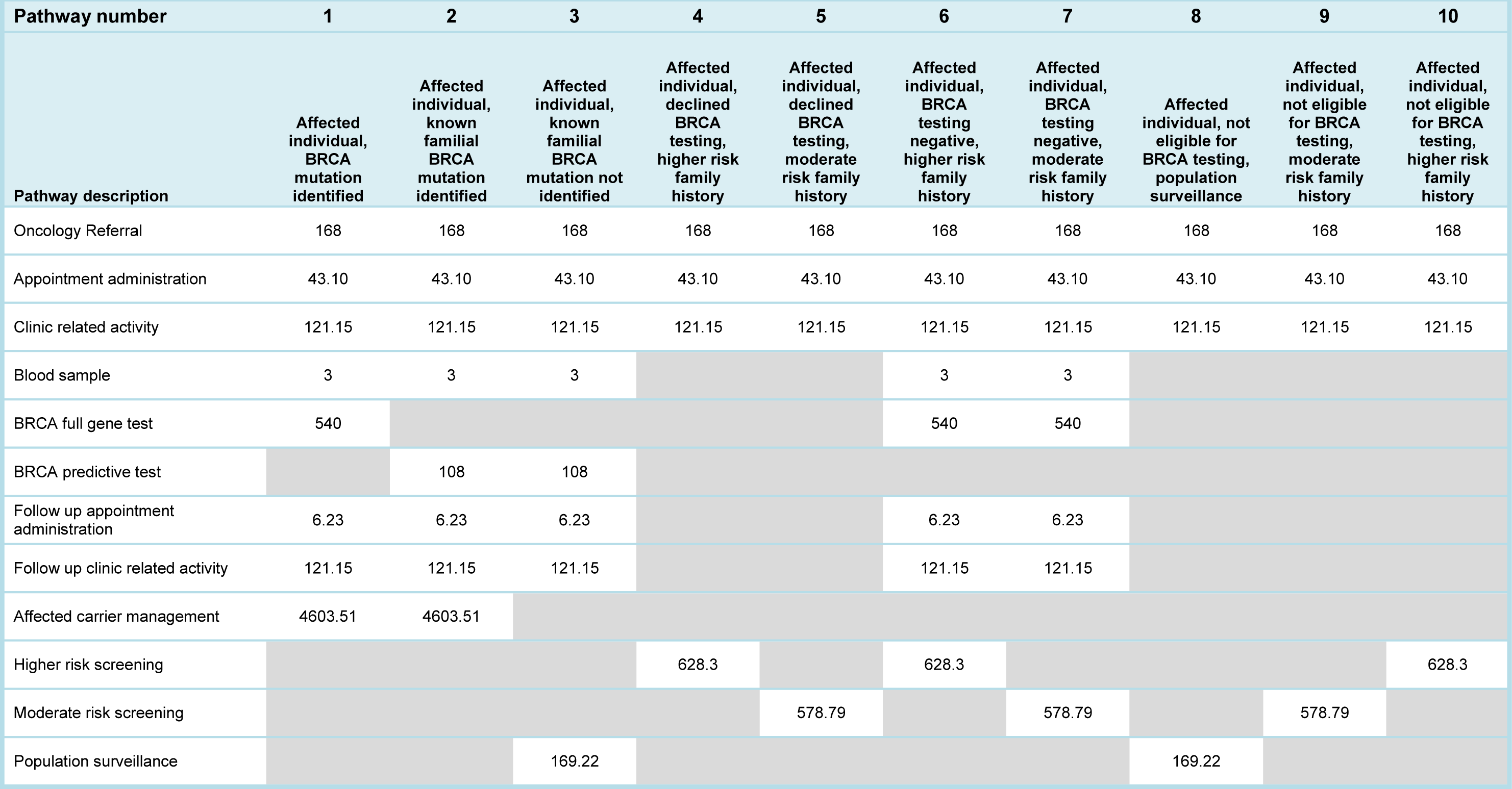
**Individuals affected with breast and/or ovarian cancer – units of activity and associated costs for each**

**Supplementary Table 2:**
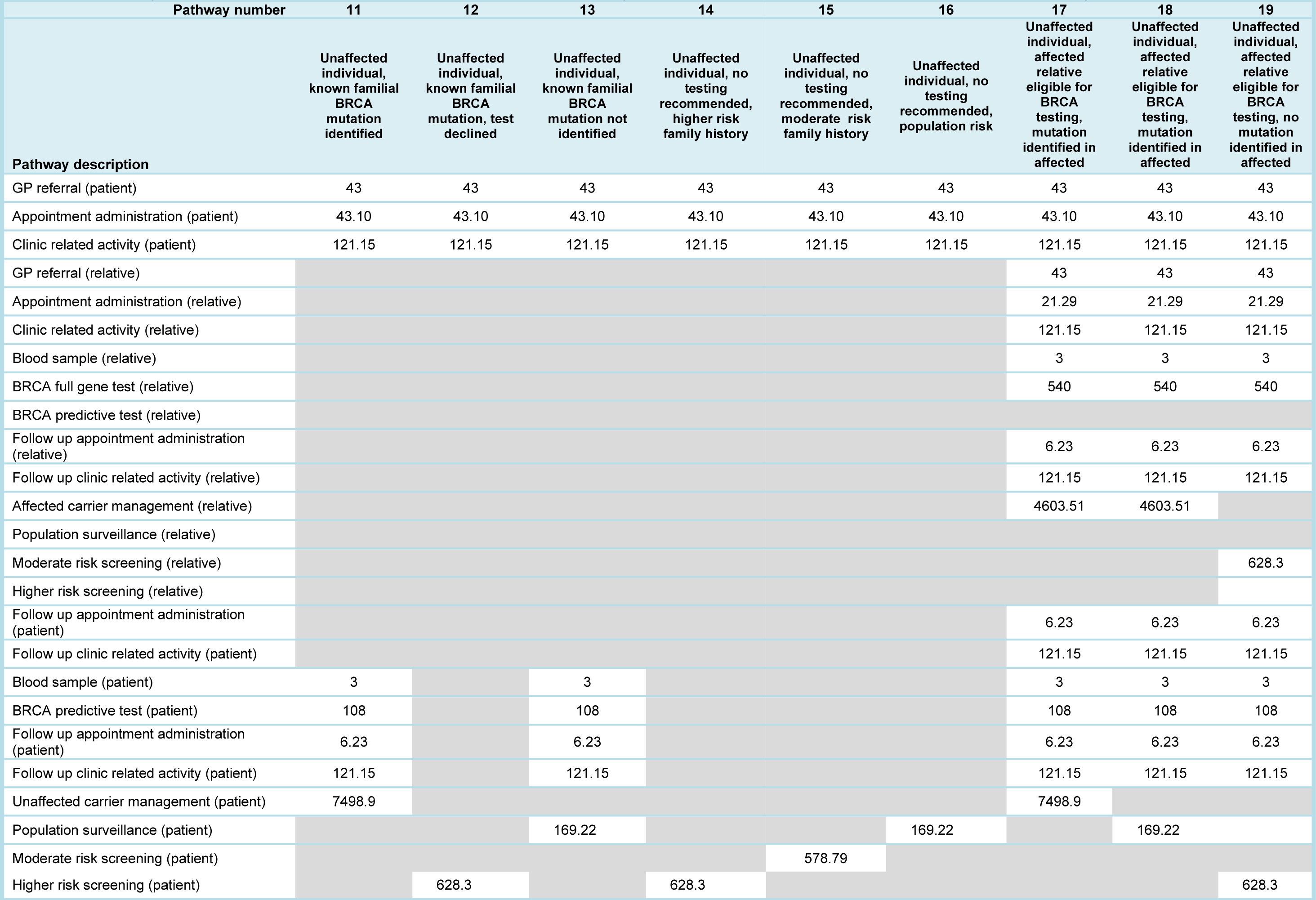
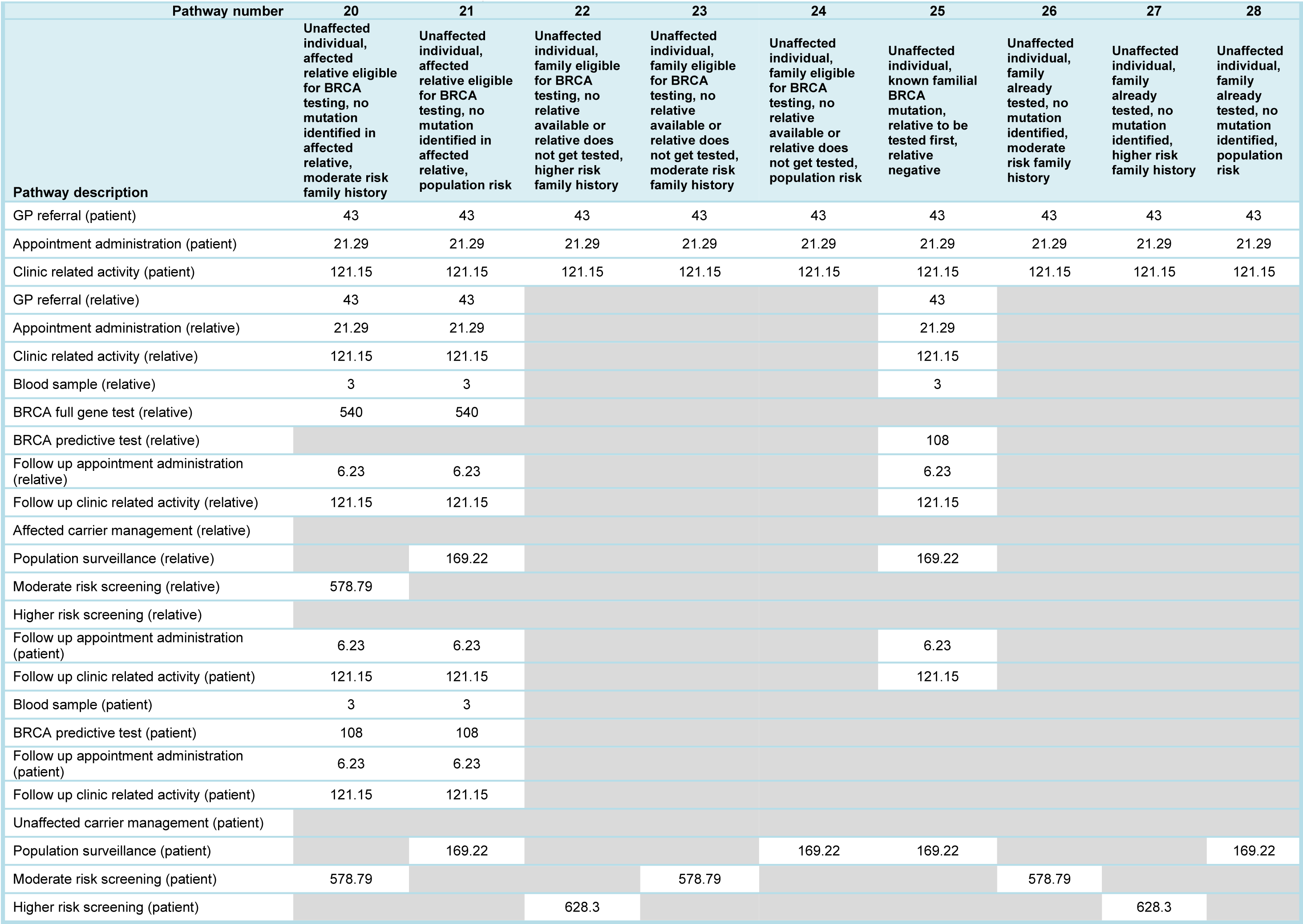
**Unaffected individuals, units of activity and associated costs for each patient pathway**

## Appendix 1

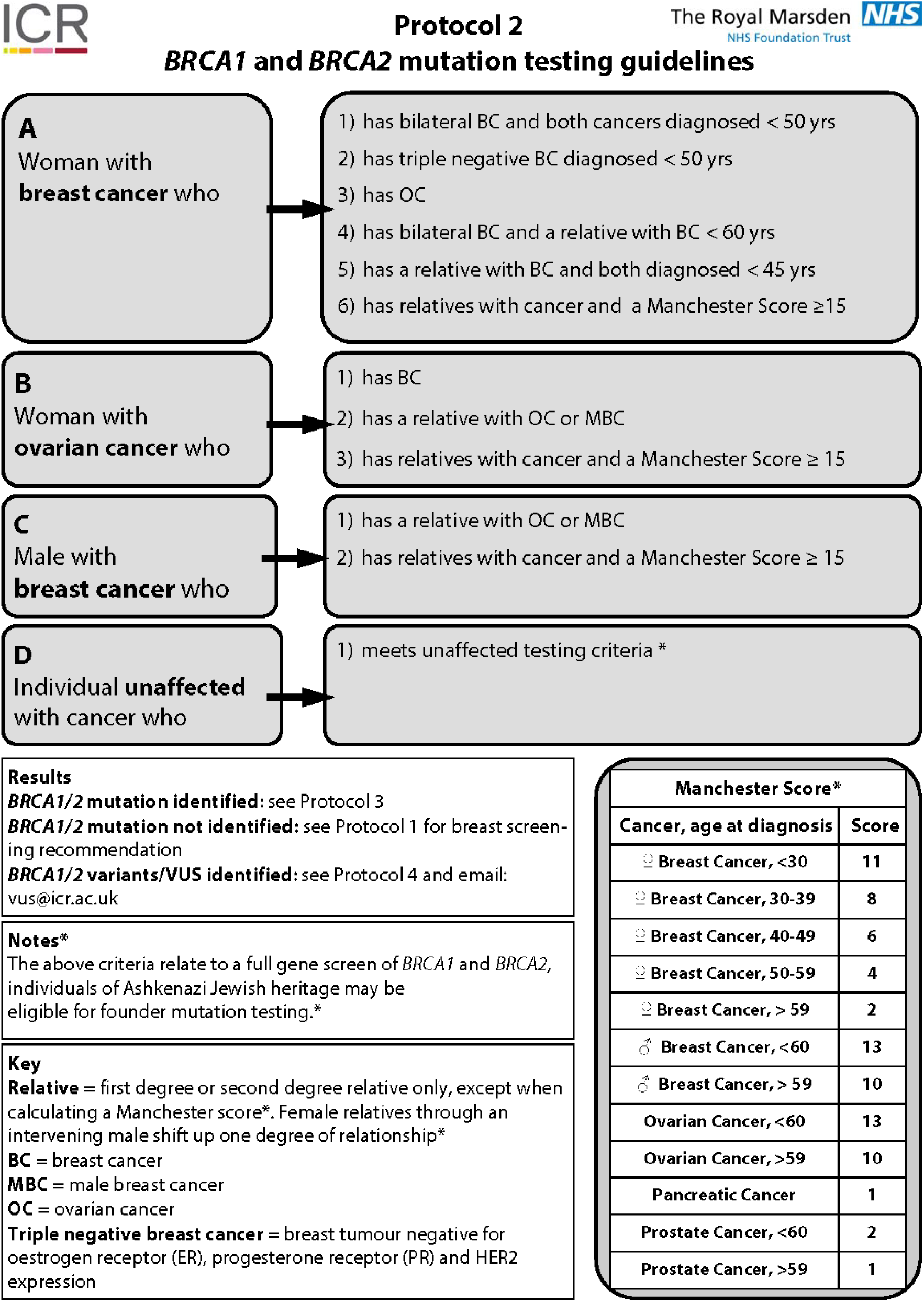

## Appendix 2

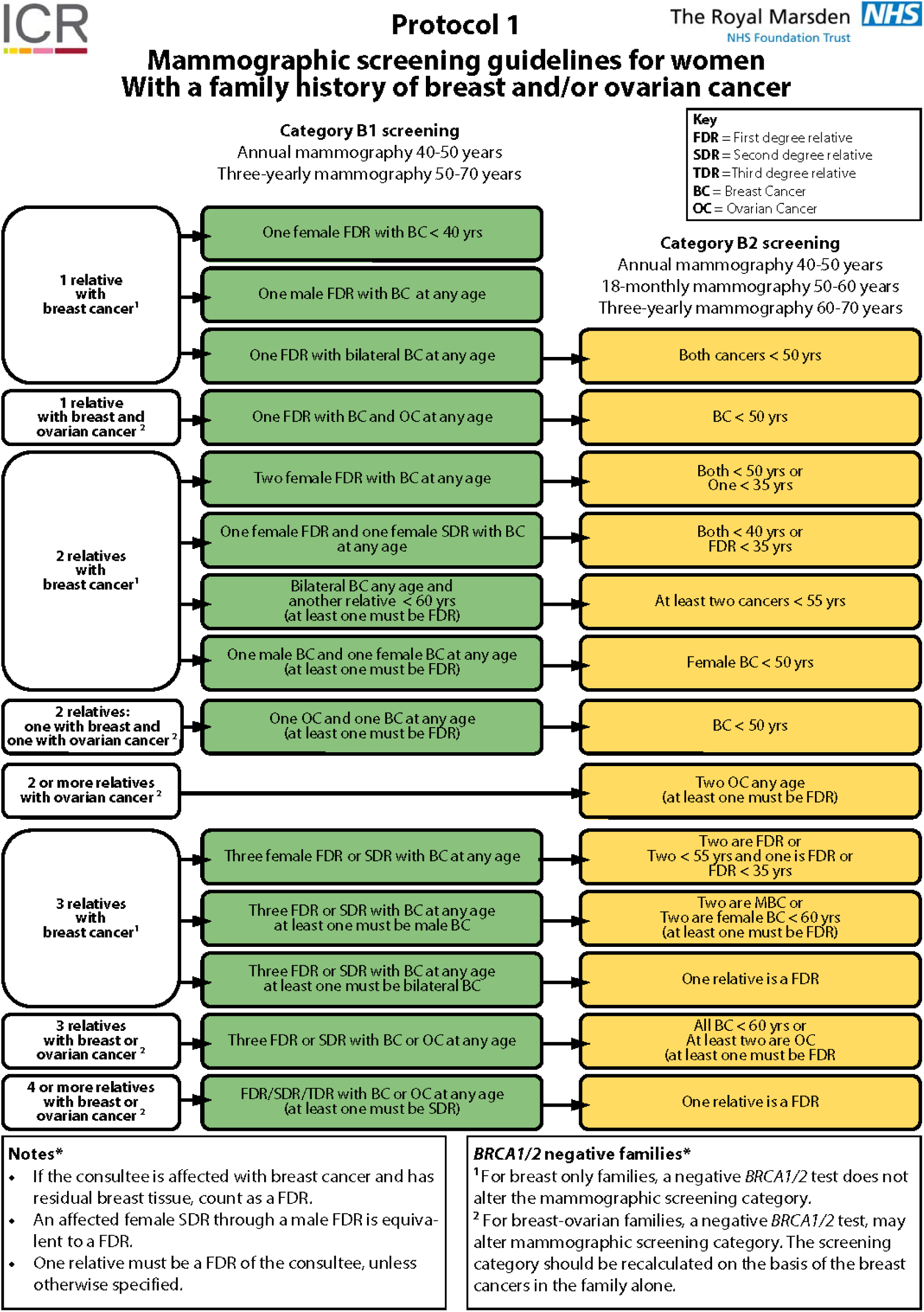

